# Why Environmental Biomarkers Work: Transcriptome-Proteome Correlations and Modeling of Multi-Stressor Experiments in the Marine Bacterium *Trichodesmium*

**DOI:** 10.1101/2021.06.22.449505

**Authors:** Nathan G. Walworth, Mak A. Saito, Michael D. Lee, Matthew R. McIlvin, Dawn M. Moran, Riss M. Kellogg, Fei-Xue Fu, David A. Hutchins, Eric A. Webb

## Abstract

Ocean microbial communities are important contributors to the global biogeochemical reactions that sustain life on Earth. The factors controlling these communities are being increasingly explored through the use of metatranscriptomic and metaproteomic environmental biomarkers, despite ongoing uncertainty about the coherence between RNA and protein signals. Using published proteomes and transcriptomes from the abundant colony-forming cyanobacterium *Trichodesmium* (strain *T. erythraeum* IMS101) grown under varying Fe and/or P limitation and/or co-limitation in low and high CO_2_, we observed robust correlations of stress induced proteins and RNAs (i.e., those involved in transport and homeostasis) that can yield useful information on nutrient status under low and/or high CO_2_. Conversely, transcriptional and translational correlations of many other central metabolism pathways exhibit broad discordance. A cellular RNA and protein production/degradation model demonstrates how biomolecules with small initial inventories, such as environmentally responsive proteins, can achieve large increases in fold-change units, as opposed to those with higher basal expression and inventory such as metabolic systems. Microbial cells, due to their close proximity to the environment, tend to show large adaptive responses to environmental stimuli in both RNA and protein that result in transcript-protein correlations. These observations and model results demonstrate a multi-omic coherence for environmental biomarkers and provide the underlying mechanism for those observations, supporting the promise for global application in detecting responses to environmental stimuli in a changing ocean.

## Introduction

A recurring question in the interpretation of transcriptome and proteome datasets is the extent to which they co-vary. Messenger RNA (mRNA) and protein levels have been reported to generally be uncorrelated within a single cell, and only modestly correlated in populations of cells, due to differences in half lives and degradation rates or phase variation within a population, respectively ^1^. It has also been shown that genes and their corresponding proteins associated with different cellular processes (e.g., core central vs stress metabolism) may retain varying degrees of correlation ^2^. While observation of correlations in larger organisms may be challenging due to internal tissues being more remote environmental signals, microbes due to their small size and immersion within the environment often maintain multiple adaptive response capabilities to common environmental stimuli. Microbial datasets with transcriptomic and proteomic methods applied to the same experiment(s) are becoming more common and hence could aid in detection and interpretation of key environmental processes. In recent years, concurrent measurements of transcripts and proteins have been conducted in marine microbes such as *Pelagibacter* ^3^, *Prochlorococcus* ^4^, the marine diatom *Thalassiosira pseudonana* ^5^, the brown alga *Aureococcus anophagefferens* ^6^, the polar alga *Phaeocystis antarctica* ^7^, and the diazotrophic cyanobacterium *Trichodesmium* ^8^. These studies also observed varying extents of transcriptome-proteome coupling, with correlations observed particularly for genes involved in responding to environmental stresses, such as P, Fe, or vitamin B_12_ limitation. Despite these efforts, there is arguably a lack of consensus in the marine ecology community regarding the extent that RNA transcripts and proteins should correlate, with many having the opinion that correlations do not occur.

In this study we synthesize the results from a number of *Trichodesmium* transcriptome-proteome datasets across a spectrum of conditions to understand their transcriptional and translational responses on a mechanistic level. *Trichodesmium spp*. are filamentous, buoyant microorganisms that can commonly grow in macroscopic colonies in close association with other microbes and have the capability to form massive blooms ^9^. Given their ability to fix both carbon and nitrogen, *Trichodesmium* spp. have relevance to both global productivity and biogeochemistry ^10,11^. Together with other marine microbes, they can impact both ecosystem stability and climate feedbacks ^12^. *Trichodesmium* is among one of several oceanic cyanobacteria that are globally significant sources of N ^10,13^, as well as unicellular forms (*Crocosphaera* spp., *Candidatus Atelocyanobacterium thalassa* and *Cyanothece*) and other heterocystous, symbiotic forms ^11^, that can have relatively high cell numbers ^14^ and biogeochemical importance ^15,16^. *Trichodesmium* is considered one of the most important diazotrophs in many tropical and subtropical regimes ^10,17^. As a result, research emphasis has been placed on developing field-ready, nutrient-limitation/stress markers to define the factors that control growth and N_2_ fixation in this important genus. These combined efforts have shown that iron (Fe) and phosphorus (P) primarily limit *Trichodesmium* across much of the global oceans ^10,11,18,19^. However, comparatively fewer studies have been conducted examining the interactions among future high CO_2_ concentrations, Fe, and P in the context of holistic, molecular physiology.

A motivation in understanding the dynamics of RNA and protein responses is the interpretation of natural microbial populations, and in particular the expression of genes that can provide clues to ecological or biogeochemical processes at play. Advances in sequencing (e.g., metagenomics/metatranscriptomics) and mass spectrometry (e.g., proteomics) technologies have enabled worldwide microbial characterizations from different biogeochemical regimes (e.g., Global Ocean Sampling Dataset ^20^). While these data have revolutionized how we think about microbially mediated processes, extrapolating biogeochemistry from relative abundances in metagenome data may result in biased interpretation. For example, it is now recognized that globally abundant microbial species initially identified via metagenomics may not necessarily be dominant members of the transcriptionally active microbial community that is primarily responsible for biogeochemical turnover ^21^. While metagenomics studies serve as powerful hypothesis-generating datasets and have unmasked previously underappreciated ecosystem processes (e.g., proteorhodopsin diversity ^20,22^), they have also underscored the need to understand the dynamics of intracellular molecular processes in the context of organismal physiology both in the laboratory and field. Thus, it is important to investigate molecular dynamics of biogeochemically-important microbes under differing nutrient and temporal regimes to resolve mRNA-protein dynamics that ultimately drive important biogeochemical processes. Here we focus primarily on N_2_ fixation. In past studies we characterized the physiological and evolutionary responses of the globally important, photoautotrophic, N_2_-fixing cyanobacterium, *Trichodesmium erythraeum* IMS101 (hereafter IMS101) to high CO_2_. Using these cell lines as starting genetic backgrounds (i.e., replicates adapted to both 380 μatm and 750 μatm CO_2_), we then conditioned them to iron-limited, phosphorus-limited, and iron/phosphorus (Fe/P) co-limitation to define their predicted climate change-impacted Fe- and P-stress molecular responses ^23^. Consistent with other studies ^24^, this demonstration of the fitness advantage conferred by Fe/P ‘balanced limitation’ compared to single limitation alone appears to contradict the long-standing Liebig limitation model and has implications for global biogeochemical cycles in both the current and future ocean such as increased exogenous nitrogen scavenging coupled to decreases in N_2_ fixation ^23,25,26^.

Here we build on and synthesize our prior results to examine steady-state transcriptional/translational/physiological relationships in the context of environmentally relevant (e.g., Fe and/or P affected) nutrient regimes using our high and low CO_2_-adapted, IMS101 cells lines ^23,25,27^. As mentioned, numerous studies utilizing diverse techniques ranging from chemical quotas, enzyme activities to gene or protein-based molecular stress markers have pointed to the importance of Fe and P to *Trichodesmium* N_2_ fixation *in situ* ^18,28,29^. Despite the value of molecular markers for indicating species specific bioavailability of nutrients, concerns persist that gene/protein expression can be decoupled with biogeochemically relevant rates like N2 fixation (e.g. ^30^) and thus give an inconsistent view of *in situ* limitation. We performed meta-analyses of our published, long-term Fe- and P-single limited and co-limited proteomic and transcriptomic datasets ^23,25^ to determine which genes and proteins give congruent expression patterns during mid-day across 8 experimental conditions vs those that do not. This analysis shows that major components of core metabolic pathway genes and proteins show incongruent responses to environmental perturbations whereas many well-classified nutrient-responsive genes/proteins show consistent correlations under both low and high CO_2_ regimes. A model of the cellular inventories for RNA and protein biomolecules that compares the timing of biosynthesis and degradation was created that provides a mechanistic explanation for the observed coherence of responses to environmental perturbations. Finally, the environmental biomarkers that have been characterized thus far in marine microbes are briefly reviewed.

## Materials and Methods

### Culturing and Molecular Extractions

Detailed culturing and protein ^23^ and RNA ^31^ extraction methods can be found in our previously published papers.

### Differential expression analysis

Differential expression data was from previously described datasets ^25^.

### Multivariate and Pairwise Analyses

Redundancy analysis was conducted on TMM-normalized transcript levels and nutrient concentrations using the vegan package ^32^. RDA ordinations were compared via the Procrustes test with the “protest” function in vegan using default settings.

Nonmetric multidimensional scaling (NMDS) was conducted using the “metaMDS” function from the vegan package with default settings (Bray-Curtis dissimilarities computed using Wisconsin double standardization) except for “autotransform=FALSE”. Hierarchical clustering with multiscale bootstrap resampling was conducted on Bray-Curtis dissimilarities calculated from TMM-normalized transcript levels using the pvclust package ^33^ with the following settings: method.dist=“euclidean”, method.hclust=“average”, nboot=1000.

The “cor.test” function in “R” was used to calculate Pearson correlation coefficients on log2 fold changes (relative to the r380 condition; see below) of average TMM-normalized transcript levels vs log2 fold changes of average normalized spectral counts of the corresponding protein product detected in Walworth et al. 2016 ^23^. Significance tests between mean correlation coefficients were carried out via the “perm” package^34^ using a two-sample permutation test with Monte Carlo simulation and 2000 permutations.

### Gene Ontology (GO) enrichment analysis

As previously described in Walworth et al. 2016 ^31^, Gene Ontology (GO) mappings for *Trichodesmium* were downloaded from the Genome2D web server (http://genome2d.molgenrug.nl). The “phyper” function in “R” (R Core Team 2014) was used to test for significant enrichment of GO pathways and p-values were corrected with the Benjamin and Hochberg method ^35^ using the “p.adjust” function (p ≤ 0.05). Finally, genes in enriched GO categories were manually checked.

### RNA-Protein Model

A model simulating the biosynthesis of RNA transcripts and proteins was created in MATLAB (MathWorks) using parameters listed in Table 1. One-minute time steps over a duration of 3 days (4320 min) within a population of 100 cells was modeled. Expression was modeled with a signal parameter that varied between 0 and 1 representing the gene being turned on or off with a simple linear ramping function. Basal signal level (s_0_) was set to 0.01, representing 1% of the population or 1 cell within 100 cells with gene expression to avoid zero denominator in fold-change estimates. In an alternate scenario representing metabolic functions in continuous use, a basal expression of 20% (0.2) was used. Rate constants for production of RNA and protein (k_RNA_, k_protein_), as well as their decay (k_-RNA_, k_-protein_) were obtained from experimental measurements in *Escherichia coli* and *Saccharomyces cerevisiae* where the degradation of each type of biomolecule was inhibited and measured ^36,37^. Change in the inventories of RNA and protein biomolecules was estimated using simple production and loss terms (Equations 1 and 2):

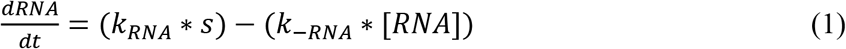

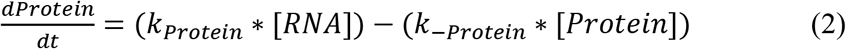

Turnover time estimates of these biomolecules are, to our knowledge, unavailable for marine organisms. While they may be different, presumably slower based on observed growth rates, it seems a reasonable assumption that the large difference between RNA and protein decay would be maintained. Hence the trends observed here are likely to be consistent or further enhanced, particularly if the RNA degradation activity in marine microbes is slower than that of *E. coli* as might be expected.

**Table 1.** Parameters in transcriptome and proteome expression model.

| Constant | Description | Value | Reference |
| --- | --- | --- | --- |
| $k_{\text{RNA}}$ | Production of RNA | 100 | |
| $k_{\text{-RNA}}$ | Degradation of RNA | $0.102 \text{ min}^{-1}$<br>(6.8 min half-life) | 1<br>(*E. coli*) |
| $d_{\text{RNA}}$ | Delay between signal and RNA | 1 min | |
| $k_{\text{protein}}$ | Production of protein | 3.5 | 3 |
| $k_{\text{-protein}}$ | Degradation of protein | $0.0017 \text{ min}^{-1}$<br>(6.9 h half-life) | 2<br>(Yeast) |
| $d_{\text{protein}}$ | Delay between RNA and protein | 180 min | |
| $S_0$ | Basal signal level 1% | 0.01 | |
| $S_1$ | Increased signal due to environmental cue | 0.2 | |
| $S_{0\text{alt}}$ | Basal signal 20% | 0.2 | |
| $S_{1\text{alt}}$ | Increased signal due to environmental cue<br>(+0.2) | 0.4 | |
| $t_{\text{start}}$,<br>$t_{\text{plateau1}}$,<br>$t_{\text{plateau2}}$, $t_{\text{end}}$ | Time of onset of signal ($t_{\text{start}}$), begin plateau<br>($t_{\text{plateau1}}$), end plateau ($t_{\text{plateau2}}$), end of decrease<br>in signal $t_{\text{end}}$; (“long slow ramp”) | 600, 650, 700, 1200<br>min | |
| $t_0$<br>$t_0$ by day | Time point denominator for fold change | 9 h (540 min)<br>540, 1980, 3420 min | |

<sup>1</sup> Selinger et al., 2003 <sup>2</sup> Pratt et al., 2002; 6.9 – 34 h protein half lives reported <sup>3</sup> Tuned using 1 minute signal to be 6-7 proteins per transcript based on Moran et al., 2013

This model has similarities to that of Moran *et al.* 2013 ^38^, but is applied here to focus on any coordination between RNA and protein and how results affect fold change estimates. Moran *et al*. instead explored random triggers for transcript production using 10 random events per day of 1 transcript per event, and then simulated environmental stimuli by increasing to 3 transcripts per event. In contrast, the model in the present study estimates production of RNA and protein molecules based on their upstream signal (signal, transcripts) as described in equations 1–2, rather than assigning a constant number of seven protein copies per transcript. Moran *et al.* observed the total number of mRNA molecules in a typical marine bacterium was only ~200 per cell^38^. These observations were incorporated into the model by allowing kRNA to be 100 to enable signal input to be transcribed into 100 transcripts across a population of 100 cells, and then tuning the protein production k_Protein_ parameter to produce 6-7 proteins per transcript also based on Moran et al., 2013. Similarly Moran et al. used a RNA half-life of 1.5 minutes, whereas this model uses a more conservative RNA half-life of 6.8 minutes based on a tiling microarray study in *E. coli*^36^

## Results and Discussion

### Global comparison of CO_2_-impacted, nutrient-limited molecular dynamics

A comprehensive depiction of the prior experimental design can be found in Fig. S1a and in Walworth et al. 2016 ^23^ in addition to physiological and proteomic results. To investigate shifts in molecular metabolism underlying Fe-limited, P-limited, and Fe/P co-limited metabolisms as they interact with predicted future CO_2_ conditions, we examined the global transcriptional output in each scenario from Walworth et al. 2018 ^8^ and their corresponding translational correlations with proteins identified in Walworth et al. 2016 ^23^. Across all treatments, we detected transcription of ≥96% and translation of 37% (see below) of the IMG annotated genes. Instances where comparisons between transcripts and proteins were not possible due to one of those biomolecules not being detected were excluded. Initial examination of the dataset in log2 fold change space revealed a cluster of points around the origin with a subset of data extending into the northeast and southwest quadrants implying a co-varying relationship between some transcript-protein pairs (Fig. 1). It is important to note that non-translated mRNAs can also potentially represent ‘irrelevant expression’, if these transcripts are not regulatory. Thus, there is much to be learned from future studies reconciling the constantly developing techniques of RNA and protein based ‘omics. Yet, there have been few studies to date comparing multi-condition proteomes with concomitantly taken transcriptomes.

**Figure 1.**
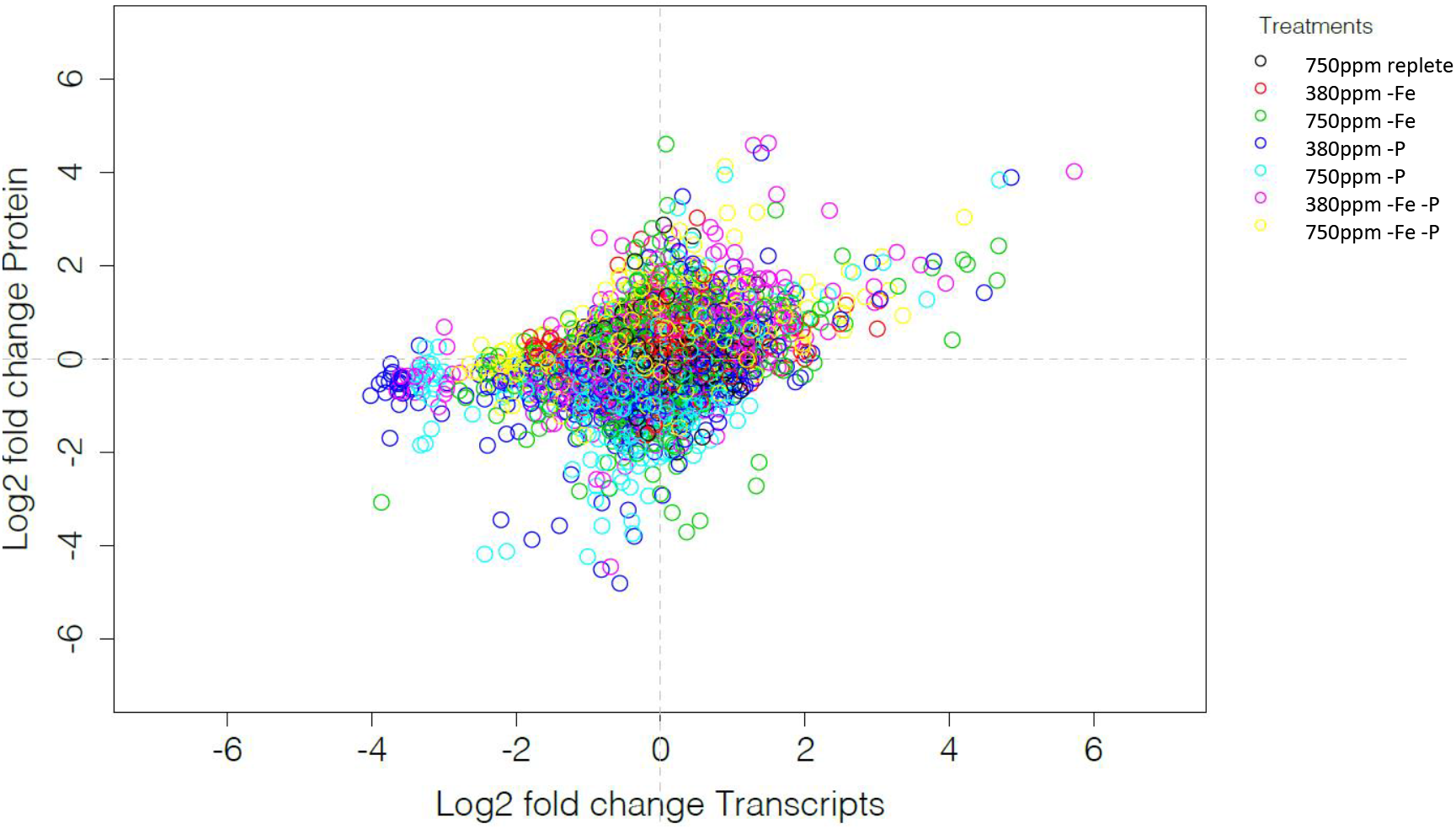
Overall comparison of proteome-transcriptome data in log2 fold space for eight experimental treatments, where treatments varied CO_2_, iron and phosphorus (see Fig. S1 for full experimental design) in cultures of the marine cyanobacterium *Trichodesmium erythraeum* IMS101. Legend shows symbols of seven treatments relative to the 380ppm CO_2_ replete treatment. Transcript data is the average of biological duplicates, protein data is the average of biological triplicates.

To this end, we first conducted exploratory analysis on the global transcriptional output without constraining for environmental variables (see below) in order to observe how replicates clustered without imparting treatment information (i.e. no assumptions). We used both nonmetric multidimensional scaling (NMDS) and hierarchical clustering with multiscale bootstrap resampling (replicates = 1,000) ^40^. NMDS revealed consistent nutrient-limited patterns across replicates (Fig. S1b), which was further supported by hierarchical clustering (Fig. S1c) that generated two high-confidence clusters with AU-bootstrap p-values > 0.95 (i.e. rejection of the null hypothesis that the clusters do not exist at the 0.05 significance level). In these unconstrained analyses, the 380-Fe condition (low-CO_2_, low iron) demonstrated greater inter-replicate transcriptional variation, although both replicates were indeed Fe-limited as evidenced by both reduced growth ^23^ and several upregulated Fe-stress genes (Fig. S1b; see below and Walworth et al. 2018 ^8^). Interestingly, Fe-limited treatments (380- and 750-Fe) clustered with the nutrient replete ones (r380 and r750) supporting the notion that P-limitation evokes a more varied molecular response than Fe-limitation relative to replete nutrient regimes (Fig. S1c). This trend was further corroborated through hierarchical clustering of transcript levels of all differentially expressed (DE) genes (i.e., those DE in any treatment relative to r380; n = 1943; Fig. 2a). To better resolve sources of variation producing these groupings, we further clustered upregulated (n = 951; Fig. 2b) and downregulated (n = 1074; Fig. 1c) transcripts, which yielded strikingly different trends among Fe-limited and replete conditions. Upregulated genes exhibited significantly different (bootstrap confidence level > 0.95) clusters among Fe-limited and CO_2_ regimes relative to those produced by clustering all DE genes while those of downregulated genes maintained similar groupings to those of DE genes. These patterns suggest that Fe-limited and replete clusters produced from all DE genes (Fig. 2a) may have been driven more so by downregulated transcriptional variation of the DE gene pool. Hence, mechanistic adaptations to high CO_2_ and responses to Fe-limitation both seem to be more similar to each other than to molecular responses under P-limitation. Interestingly, both up- and downregulated P-limited clusters were statistically analogous to those produced by clustering all DE genes (Fig. 2a) and total genes (Fig. S1c), which further corroborates the notion that P-limitation evokes a highly conserved yet more varied transcriptional rearrangement than either high CO_2_ or Fe-limitation.

**Figure 2.**
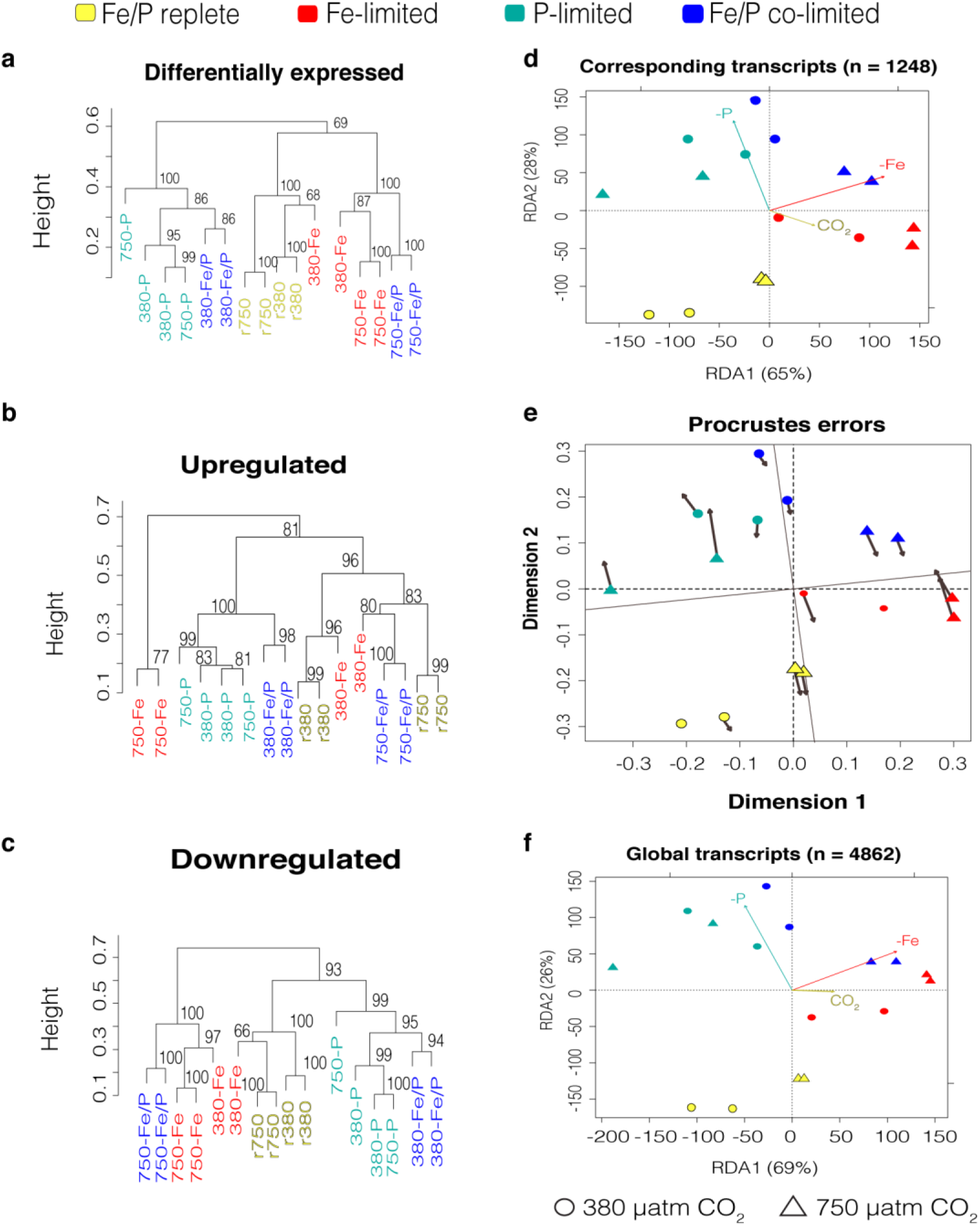
Global molecular analysis. (a) The top panel shows a redundancy (RDA) analysis of normalized transcript abundances of genes whose protein products were also detected in Walworth et al. (2016a). The middle panel shows a Procrustes analysis between corresponding-RNA-RDA and prot-RDA ordinations indicating close agreement evidenced by small vector residuals (r = 0.75; P < 0.001). The bottom panel displays the same redundancy analysis as in the top panel except now including all detected gene transcripts. (b) Hierarchical clustering of Bray-Curtis dissimilarities with multiscale bootstrap resampling calculated from normalized transcripts of differentially expressed (top panel), upregulated (bottom left), and downregulated (bottom right) genes. Numbers at dendrogram nodes are approximately unbiased p-values calculated from multiscale bootstrap resampling.

Next, we employed constrained multivariate methods that attempts to explain variation based on the given environmental variables (e.g. RDA analysis) for both transcripts and proteins. When relating nutrient-limited proteome variation identified via redundancy analysis (RDA) in Walworth et al. 2016 ^23^ (n = 1248; 37% of genome coding potential) to that of their corresponding gene transcripts (corresponding-RNA-RDA; Fig. 2d), similar nutrient-limited patterns across replicates were observed with most of the proportional variance explained by the environmental variables for both transcripts (adjusted R^2^ = 0.56) and proteins (adjusted R^2^ = 0.59), respectively. Moreover, a statistically significant correlation was observed (r = 0.75; P < 0.001) when comparing the corresponding-RNA-RDA and prot-RDA ordinations via permutational, least-squares orthogonal analysis (i.e., Procrustes analysis; Methods) indicating that, overall relative transcriptional variation among treatments was captured in the corresponding proteome variation (Fig. 2e).

Performing the same RDA analysis on the global transcriptional output (global-RNA-RDA; Fig. 2f; adjusted R^2^ = 0.50; n = 4862) displayed a strikingly consistent ordination relative to that of both the prot-RDA (represented in Figure 1c in Walworth et al. 2016^23^ and corresponding-RNA-RDA (Fig. 2d). In fact, when analyzing congruence of the global-RNA-RDA to both the prot-RDA ^23^ and the corresponding-RNA-RDA, respectively, a robust, statistically significant correlation was observed relative to both the prot-RDA (r = 0.75, P < 0.001) and the correlated-RNA-RDA (r = 0.98; P < 0.001). Taken together, these analyses indicate that both transcriptional and translational variation inherent in the overlapping 37% of the detected proteome and 96% of the detected transcriptome are not only representative of one another across nutrient-limited conditions, but also of global transcriptional variation. In other words, this congruence implicates that the detected proteome variation is indeed representative of overall treatment-specific, transcriptional variation in IMS101 for Fe, P, Fe/P, and CO_2_.

Another striking observation is that 380-Fe/P replicates were more closely associated to P-limited treatments (380- and 750-P) while 750-Fe/P replicates were more correlated with Fe-limited ones (380- and 750-Fe) for both ordination-based (RDA, Fig 2d,f; NMDS, Fig. S1b) and clustering analyses (Fig. 2a,b,c and Fig. S1c). Hence, these trends imply that regardless of the amount of variation explained by the given environmental variables, the global transcriptional profile of Fe/P co-limited metabolism is more similar to that of P-limitation when adapted to current CO_2_ levels (380-Fe/P) but is more similar to Fe-limitation when adapted to high CO_2_ (750-Fe/P).

### From global to pairwise mRNA-protein correlations

Bacterial messenger RNA (mRNA) typically have much shorter half-lives than those of proteins and can be rapidly transcribed in response to quickly changing environmental conditions ^41^. For example, in an *E. coli* cell, many mRNAs are typically degraded within minutes, whereas their corresponding proteins can have longer lifetimes than the cell cycle ^42^. These data indicate that intracellular mRNA copy number at any instant may typically reflect recent transcriptional activity, while protein levels in the same instant can represent a relatively longer history of accumulated expression. Indeed, simultaneous measurements of mRNA and their corresponding protein concentrations have been shown to be uncorrelated within a single cell at a single time point, while mRNA levels integrated over many cells in a population generally correlate with protein concentrations ^42^. Different environmental variables invoking specific metabolic pathways can also impact overall mRNA-protein correlations ^1,2^.

Unlike prior other microbial studies investigating mRNA-protein correlations across one or two treatments ^2^, we examined correlations across eight different growth conditions. As in most organisms to date, IMS101 transcript levels modestly correlated to their corresponding protein abundances across all treatments (n = 1,246; squared Pearson correlation coefficient, r^2^ = 0.36), suggesting that other forms of regulation might need to be invoked to explain the majority of variation ^1^. Upon calculating Pearson correlation coefficients across all treatments for genes that were differentially expressed (DE) in at least one experimental condition relative to r380 versus those that were not called as differentially expressed (NDE) in any condition, we observed a significantly greater mean correlation coefficient for DE (r^2^ = 0.35) vs NDE (r = 0.03) genes (p < 0.0005; two-sample permutation test with Monte Carlo simulation). This same robust trend was also observed when comparing average correlation coefficients for DE vs. NDE genes for each pairwise treatment comparison (Fig. S2). Hence, this greater correlation suggests that transcription and translation of environmentally responsive genes and their protein products may be more closely coupled on average than those that do not respond to changing environmental conditions.

### Pathway-specific molecular relationships

To search for metabolic pathways harboring highly correlated transcript and protein abundances, we calculated Pearson’s correlation coefficients (PC) and kept genes retaining PC values ≥ 0.7 and p-value < 0.05. We then mapped genes onto their Gene Ontology (GO) pathways and searched for significant GO enrichment using the hypergeometric test with Benjamini-Hochberg correction (FDR ≤ 0.05; Dataset S1). Enriched GO pathways included translation, nitrogen fixation, oxidation-reduction, calcium ion binding, proteolysis, iron-sulfur cluster binding, and photosynthetic electron transport/light reactions. Hence, these data suggest that biosynthesis of these protein products increases with increasing transcript accumulation. Therefore, their abundances may be more representative of instantaneous cellular activity rather than those with low PC values, in which the latter may be a product of constitutive activity and/or reduced protein degradation. For example, well-characterized Fe- and P-stress genes exhibited high PC values in conjunction with genes involved in photosynthetic light reactions (Fig. 3). In all cases, the interaction of high CO_2_ with limiting Fe irrespective of fluctuating P (750-Fe and 750-Fe/P) yields increases in both transcript and protein levels for all Fe-stress genes relative to their low CO_2_/low Fe conditions (i.e. 380-Fe and 380-Fe/P). Hence, the expression of these Fe-stress biomarkers increases under limiting Fe in future ocean conditions suggesting their profiles to be robust for *in situ* profiling. The genes involved in photosynthesis light reactions (Fig. 3c) exhibit parallel trends possibly suggesting co-regulation by iron limitation or growth rate. Interestingly, P-stress transcript and protein levels robustly respond to limiting P irrespective of Fe and CO_2_, but increased CO_2_ has little to no effect on P-stress molecular machinery (Fig. 3b). Additionally, these data suggest that different subunits of the same phosphate transport system (i.e. Pst) could respond differently to fluctuating P as transcript and protein levels of, for example, the PstB subunit increased more considerably than those of PstS under limiting P (Fig. 3b).

**Figure 3.**
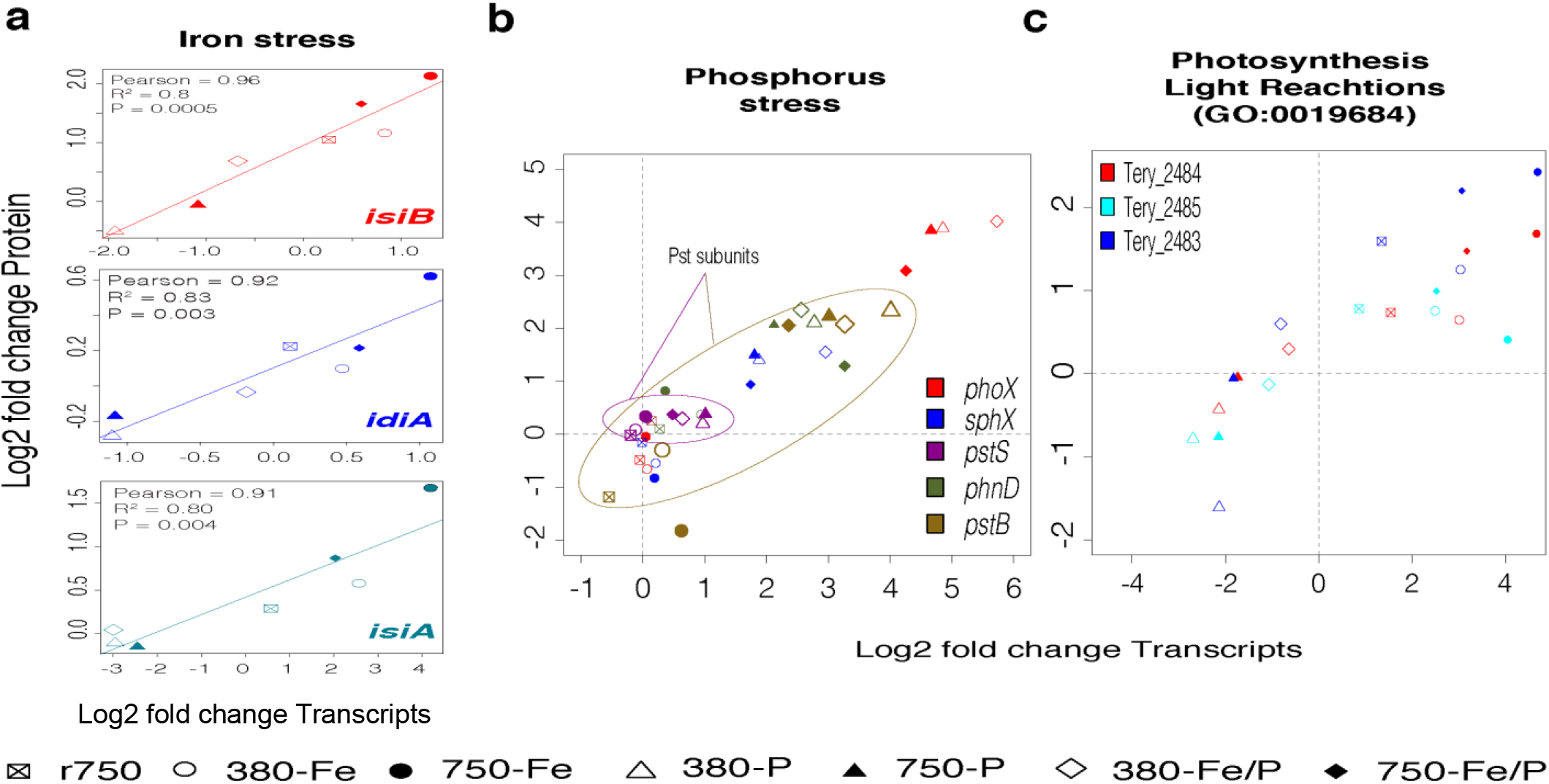
Scatterplots of log2 fold changes of transcript and protein abundances for select, highly correlated genes. (a) Shown are log2 fold change of normalized protein (y-axis) and transcript (x-axis) abundances for Fe-stress genes (b) Shown are log2 fold change of normalized protein (y-axis) and transcript (x-axis) abundances for P-stress gene. Ellipses denote different subunits of the Pst transport system. (c) Shown are log2 fold change of normalized protein (y-axis) and transcript (x-axis) abundances for photosynthesis light reaction genes. Different symbols denote different treatments.

Conversely, GO-pathways specific to the NDE pool with low PC values involve general biosynthetic processes, magnesium ion binding, proteolysis, respiration (e.g., TCA cycle), nucleotide metabolism, inosine monophosphate (IMP) biosynthesis, and amino acid metabolism (Fig. 4). Of note, decoupling of TCA gene expression and enzyme abundance in yeast has also been previously observed ^2^. Interestingly, the GO pathway assigned to photosystem I reaction center (GO: 0003989) experience significant enrichment of genes with low PC values (Dataset S1) while conversely, the overall photosynthesis light reaction GO pathway (GO:0019684) was enriched with genes with high PC values (Fig. 3c). Hence, it is important to distinguish between potentially constitutive vs. responsive components of large, multifaceted pathways with many components that can have widely differing roles (e.g., energy flow vs. structure). Overall, it seems the persistent activity of certain core pathways to maintain basic cellular homeostasis, irrespective of changing environmental conditions, may result in more decoupling of transcription and translation on average (Fig. 4), relative to pathways that synthesize new transcripts to generate protein products specific to particular environmental triggers (e.g., Fig. 3).

**Figure 4.**
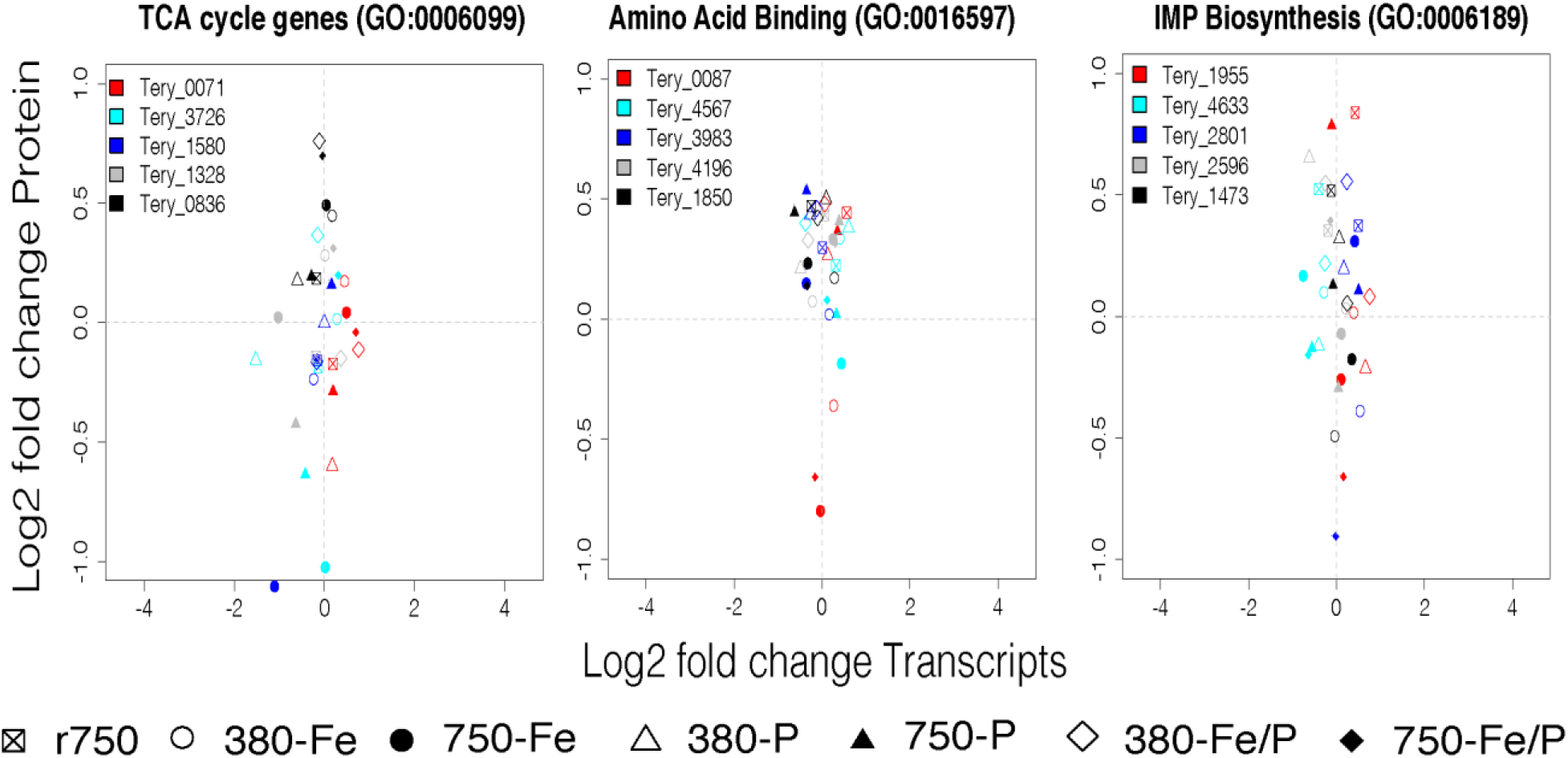
Scatterplots of log2 fold changes of transcript and protein abundances for select core metabolism genes. Shown are log2 fold change of normalized protein (y-axis) and transcript (x-axis) abundances for TCA cycle, amino acid binding, and IMP biosynthesis genes. Different symbols denote different treatments.

### RNA-Protein Modeling: Influence of biomolecule inventories on fold-change dynamics

In order to explore the potential causality of RNA-protein coupling and decoupling we generated a model for production and decay of both biomolecules (Fig. 5). Model parameters include previously published values for RNA and protein degradation based on studies of whole transcriptome and proteome degradation measurements from model organisms (*E. coli* ^36^ and yeast ^37^, respectively) as well as delays between RNA and protein expression observed in *Prochlorococcus* ^4^ (Table 1). Notably, RNA decay rates are approximately two orders of magnitude higher (~60-275 fold) than that of protein degradation. While differences in degradation rates occur between different specific transcripts and protein molecules, there is clearly a large difference between transcript and protein degradation rates, as expected based on the transient information transmission role of RNA versus the long-term functional roles of proteins. The choice of *E. coli* and yeast as organisms for obtaining decay parameters will likely vary from that of various marine microbes, with both degradation processes likely being slower within slower growing marine microbes such as *Trichodesmium*. However, the large difference in cellular decay rates of RNA and protein rates is almost certainly a universal phenomenon, and hence we expect the general trends of temporal overlap between transcripts and protein inventories modeled here to be relevant even if future taxon-specific activities vary somewhat from those used here. In particular, if the RNA degradation constant is lower in slower growing organisms due to lower abundance and/or efficiency of RNases, an increased temporal overlap between transcripts and proteins would result in greater opportunity for correlation between the two types of biomolecules. Additional parameters in the model include production of RNA and protein, as well as constants for temporal offsets between signal transduction and transcription (d_RNA_), and transcription and translation processes (d_protein_). For the latter, 3h was used, similar to the range observed in *Prochlorococcus* ^4^. Finally, the model includes initial condition parameters whose influence can be explored including basal expression levels (1% and 20%) and time settings for onset, duration, and decay of signals to initiate transcription. The notion of the transcriptional signal is generalized here, whether they occur from riboswitches, two component regulatory systems, sigma factors or other mechanisms. Moreover, the onset and decay of the regulatory signaling is also idealized here as a linear ramp (using a “long slow ramp” that repeats daily), although it could be further qualified to have positive feedbacks to increase sensory capabilities such as observed in increased expression of two component regulatory systems, e.g., phosphate regulatory systems ^43^. The influences of post-translational modifications (PTMs) were purposely not included in this model in order to allow an examination of potential range of differences in RNA and protein fold-change expression levels prior to often invoked additional PTM regulatory controls.

**Figure 5.**
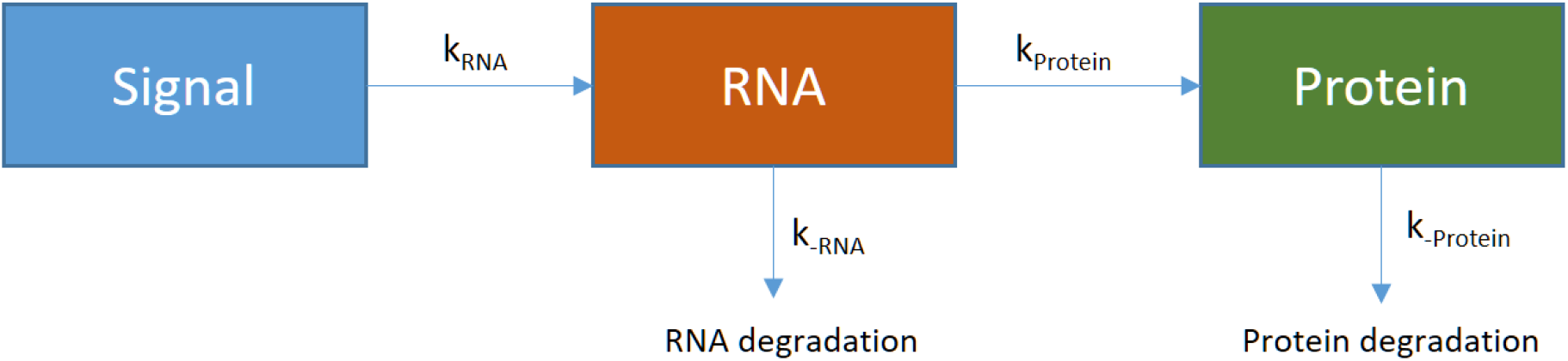
Representation of transcriptome-proteome model. Parameters described in Table 1.

The RNA-Protein expression model results imply that changes in basal expression level can have a major influence on the extent of RNA-proteome coupling and decoupling observed experimentally. For example, when there is no significant basal expression (1% or s_0_=0.01; where a non-zero value was chosen to allow fold-change ratio estimate) as would be expected prior to an environmental stimulus such as phosphate or iron stress, both RNA and protein signals respond with a large increase in inventory and fold-change (Fig. 6 and 7 left panels). In contrast, when there was a 20% basal level of expression (s_0_=0.2), as expected for routinely used metabolic machinery, a significant inventory of protein product accumulates (Fig. 6 right panels), causing the fold change signal to become muted (Fig. 6). These results are also displayed as a movie format (Supplemental Movie M1) to display the temporal unfolding of these transcript-protein inventories and their influence on fold changes results. The fold change decrease is accentuated if calculated each day as the protein inventory accumulates (e.g., the choice of what to normalize expression to in the fold change estimate). Notably, proteins that respond to environmental stress are consistent with the lower basal expression model that yields correlations between RNA and protein. In *Trichodesmium*, photosynthetic transcripts and proteins were also observed to be correlated (Fig 4c). While generally considered metabolic proteins with high inventories, this observed correlation can be particularly driven by iron limitation scenarios, which can remodel overall machinery by reducing the iron-rich photosystem I (PSI) in favor of PSII ^23^. Iron stress transcripts and proteins like *isiA* can also concurrently increase in abundance solely in times of enhanced iron stress to form PSI-IsiA-chlorophyll– protein–antenna super-complexes ^44^ while in other conditions like phosphorus stress, they show little correlation due to the lack of an environmental trigger. Additionally, diazotrophs have been shown to have strong diel cycles that affect both transcripts and proteins of photosynthetic and nitrogen fixation enzymes in order to promote intracellular oxygen protection and iron conservation efforts ^45,46^. Hence, the choice of when to sample transcriptomic and proteomic experiments can also be consequential. The short-reported half-lives of RNA molecules cause a rapid decrease in transcripts if they are not continually being produced ^38^. The expected large two order of magnitude difference in half lives of RNA and protein molecules results in a counterclockwise progression on RNA and protein space plots commonly used for assessing coherence between these biomolecules (Fig. 8). Choice of sampling time is typically estimated to allow sufficient time for RNA and protein biosynthesis, where if too early there is insufficient time for protein synthesis, and if too late RNA transcripts may have begun degradation ^7^. Notably, the magnitude of these counterclockwise traces is affected by the basal expression level, where low basal expression creates an easily observable pattern due to the high fold change values for transcripts and proteins, while high basal expression results in the circular trace to be compressed near the origin. This is indeed what many experimental datasets look like with metabolic proteins found near the origin in log2 transcript-protein space (Fig. 1), while genes that respond strongly to environmental cues have stronger signals that project into the northeast quadrant of these plots. The model results here may be overly conservative, as it seems likely that marine microbes with much slower reproduction rates than *E. coli* or yeast would also have lower concentrations of RNases and concurrent longer residence times of RNA, as mentioned above. If lower RNase activity were common among marine microbes like *Trichodesmium*, the overlapping time window for co-existence of RNA and proteins would be enhanced.

**Figure 6.**
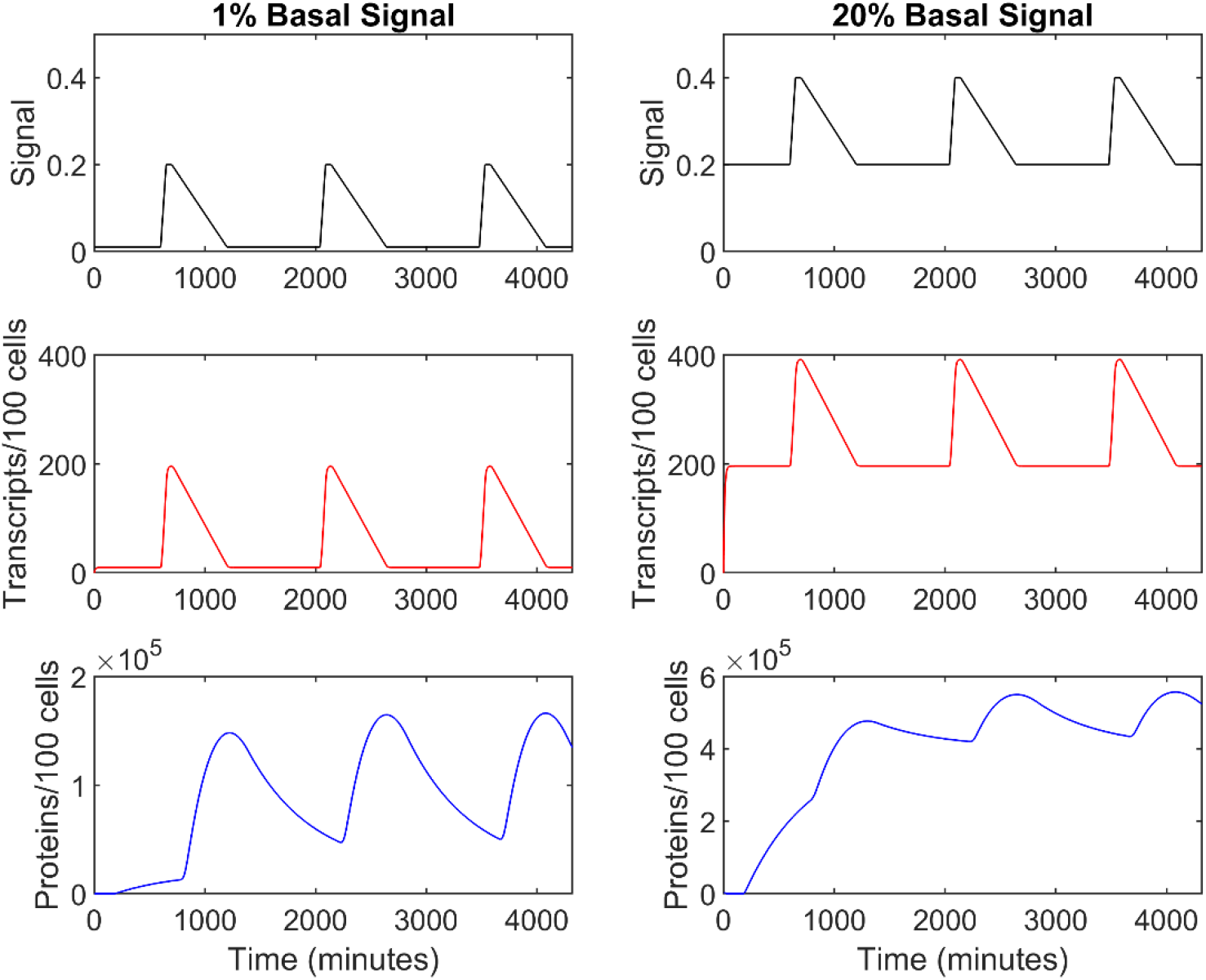
RNA-Protein expression model results. Low basal expression of 1% (left panels) and moderate basal expression (right panels), with a simulated daily expression pattern. Short and long half-lives of RNA and protein, respectively, result different inventories over time. When basal expression is increased to 20% protein inventories begin to accumulate with between successive daily cycles.

**Figure 7.**
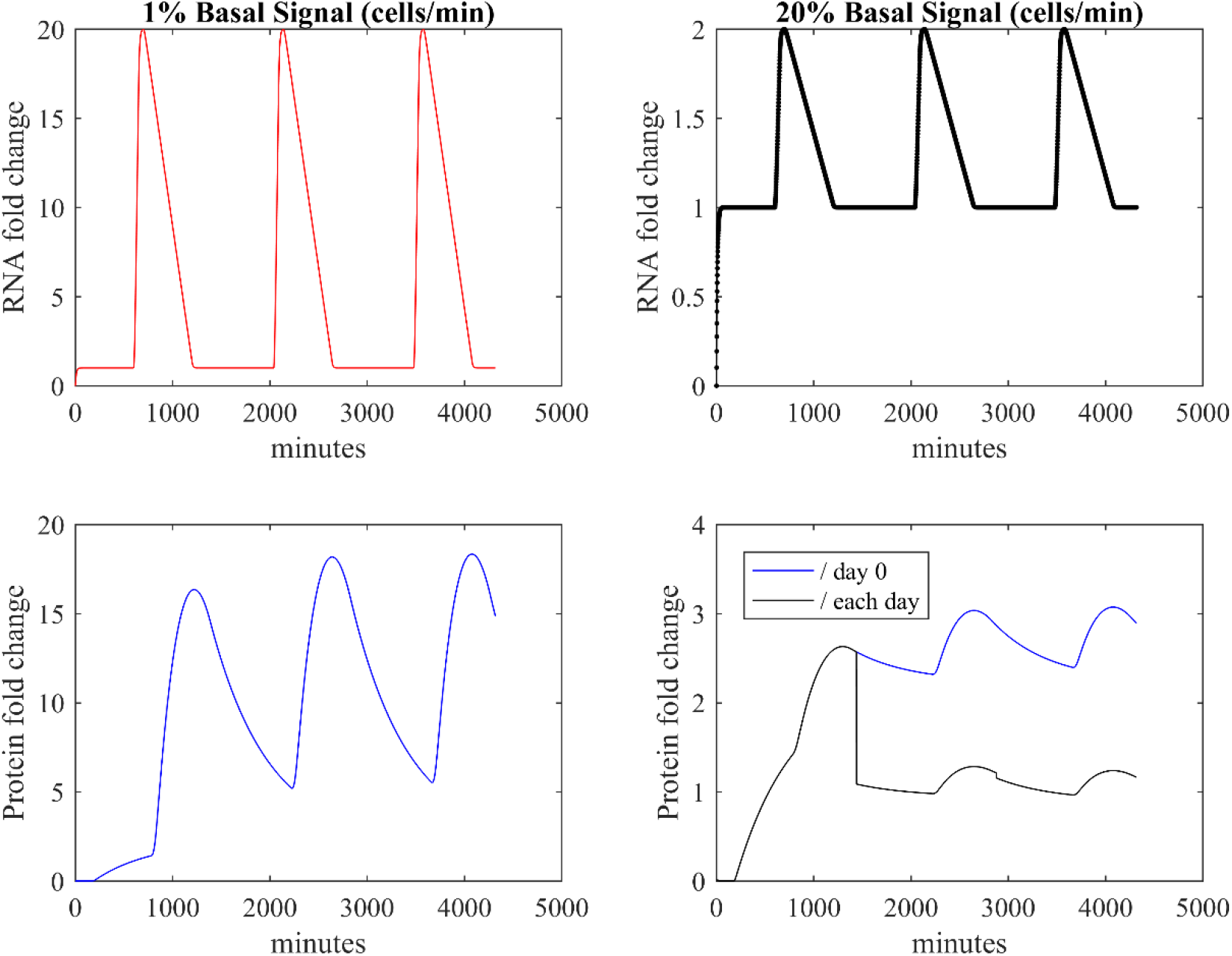
RNA and protein expression in fold change space. Model output from Figure 6 converted to fold-change space results in high values with low basal expression (1%; left panels) and muted values with some basal expression (20%; right panels). Fold-change was calculated relative to day 0 or each successive day in right panels.

**Figure 8.**
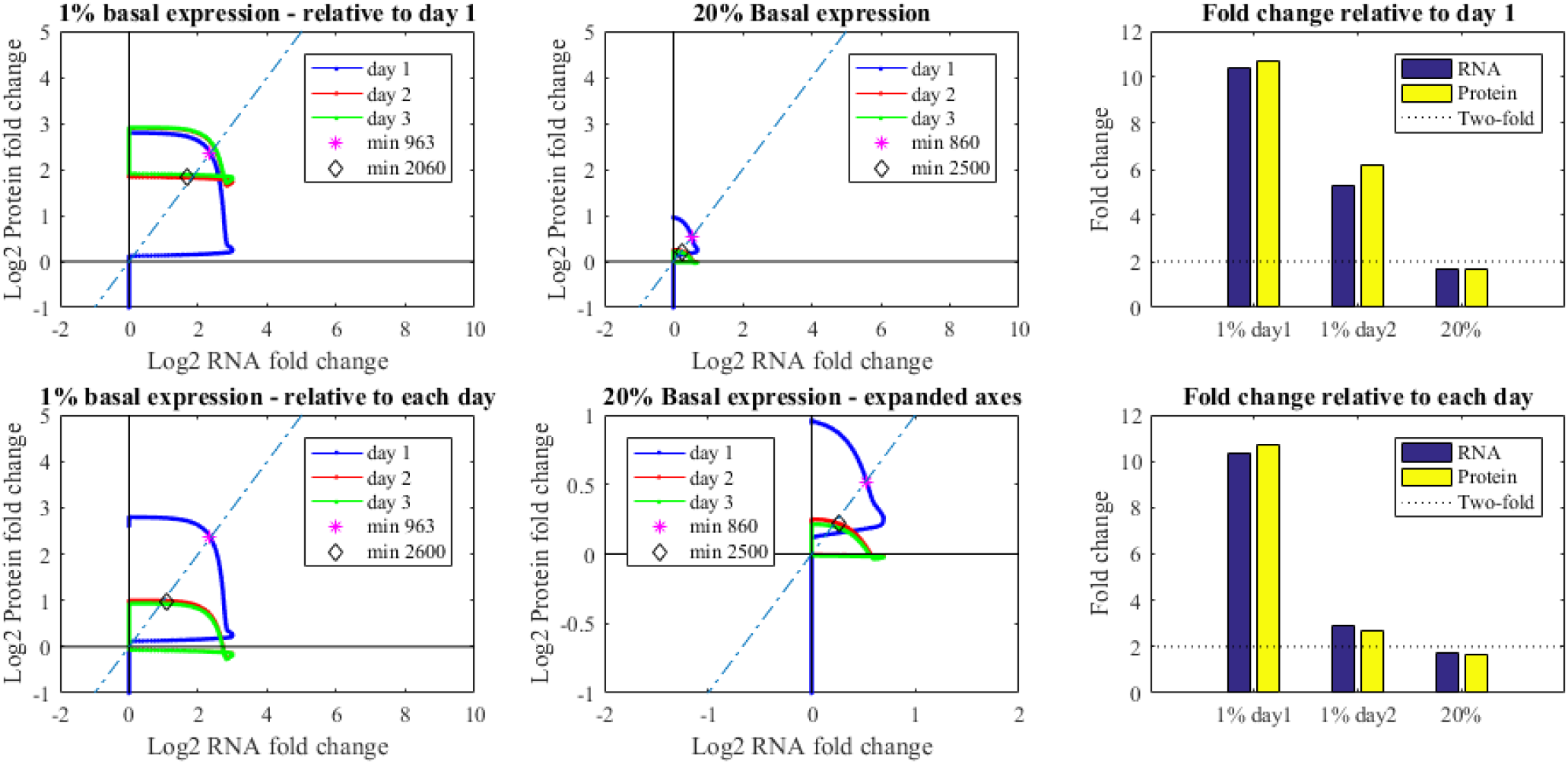
Expression patterns in transcriptome-proteome space. The succession of RNA and protein biomolecules resulted in a counterclockwise path, with more pronounced fold change signals with low basal expression (left panels) compared to higher basal expression (center panels) where the signal circular progression falls close to the origin (top middle). Coherence between biomolecules was observed as in the northeast or southwest quadrants (dotted line). The resulting maximum fold change (right panels) observed was high with low basal expression (top right) and when relative to day 1, rather than successive days as the protein inventory accumulates (Figure 6). Environmental biomarkers tend to have lower basal expression and are expressed based on regulation by environmental cues.

Based on this simple modeling approach, the (often negative) conclusions about the coherence of RNA and protein expression in microbes appear to have been clouded by use of relative units of fold change that discriminate against metabolic proteins with larger inventories. In these cases, there may be highly abundant proteins that change in inventory by 20% for example, which would be a large change in copy number, but a small change in fold change ratios. Use of absolute units for both transcripts and proteins and efforts to calibrate with standards to achieve copy number per cell estimates would resolve this problem. Importantly however for environmental applications, the successful use of environmentally responsive genes in both RNA and protein space can be explained by their large signals relative to basal expression that allows easy detection of large fold changes. In short, the lack of observed correlations of metabolic systems represented in transcripts and proteins can be explained, at least partially, by an over-reliance on ratios of data, and those observations should not detract from the statistically significant correlations of environmentally response genes in RNA-protein space and their application as research and monitoring tools in environmental microbiology and biogeochemistry.

Together these empirical observations and model results validate the utility and provide a theoretical basis behind proteins that respond to environmental stimuli in marine microbes. Because organisms that comprise natural ecosystems are frequently limited in their productivity by nutritional elements, they often have evolved specific adaptive responses to nutritional scarcity. As such, these proteins and their mRNA transcripts represent ideal environmental biomarkers to detect controls on ecosystem productivity. Biomarkers for nutritional stress in marine phytoplankton and bacteria that have been identified are shown in Table 2, including those for nitrogen ^43,48–50^, phosphorus ^5–6,51–53^, iron ^18–19, 54–61^, vitamin B_12_^62,63^, and zinc^64^. Notably, proteins involved in adaptive responses in marine microbes are often distinct from those model organisms because of their very different environmental conditions. As a result, several of these marine biomarkers have only recently been discovered and it is almost certain that more remain to be discovered. In some cases, biomarkers for required elements such as cobalt do not appear to have adaptive responses (or their chemical species they respond to has not yet been identified)^65^ and corresponding biomarkers perhaps due to the nutritional burden of such systems in highly streamlined genomes. With the natural environmental undergoing rapid and unprecedented changes, the ability to observe and understand how the oceans are changing on a global scale is possible through the application of environmental biomarkers. Demonstration that RNA and protein do correlate for environmentally responsive genes and the theory behind when and why these correlations occur is valuable in increasing confidence for study of metaproteomic and other meta-‘omic studies to the global oceans ^66–70^.

**Table 2.** Example Biomarkers for Nutrient Stress in Marine Microbes.

| **Protein Name** | **Nutrient** | **Relevant Taxa** | **Example Refs** |
| --- | --- | --- | --- |
| Urea Transporter | Nitrogen | Cyanobacteria | 43, 48-50 |
| NtcA and P-II N Regulatory Proteins | Nitrogen | Cyanobacteria | 43, 48, 66 |
| NtrX Regulatory Protein | Nitrogen | Pelagibacter | 49 |
| Ammonia transporter | Nitrogen | Cyanobacteria, Diatoms | 43, 50 |
| Alkaline Phosphatase, PhoA | Phosphorus | Cyanobacteria, Diatoms | 5, 6, 51 |
| Phosphate Transporter, PstS | Phosphorus | Cyanobacteria | 51,52,53 |
| Iron Transporter IdiA/FutA | Iron | Cyanobacteria | 18, 19, 58, 59, 60 |
| Flavodoxin IsiB and Ferredoxin | Iron | Cyanobacteria, Eukaryotic Algae | 57, 43, 7 |
| Iron Transporter SfuC | Iron | *Pelagibacter* | 61 |
| ISIP2 | Iron | Eukaryotic Algae | 7, 54, 55 |
| ISIP3 | Iron | Eukaryotic Algae | 7, 54 |
| B<sub>12</sub> transporter CBA1 | Vitamin B<sub>12</sub> | Eukaryotic Algae | 62-63 |
| Methioine Synthase MetE and MetH | Vitamin B<sub>12</sub> | Eukaryotic Algae | 62-63 |
| Zn chaperone ZCRP-A | Zinc, Cobalt | Eukaryotic Algae | 64 |
| Zn membrane protein ZCRP-B | Zinc, Cobalt | Eukaryotic Algae | 64 |
| Unknown | Co, Ni, Mn, vitamins, etc. |  | 65 |

## Conclusions

In summary, these data elucidate broad, nutrient-limited mRNA-protein dynamics following long-term high CO_2_ adaptation under multiple nutrient limitation regimes in a biogeochemically-important organism. Overall, nutrient-limited proteome variation representing 37% of the IMS101 coding potential was congruent with that of its corresponding transcripts, as well as with global transcriptional variation. Pairwise mRNA-protein transcript levels were modestly correlated to protein abundances, yet environmentally responsive transcripts were significantly more correlated than those that did not respond to any of the 8 conditions. Well-characterized stress mRNA and protein biomarkers responded robustly to their respective limiting nutrients following long-term CO_2_ adaptation, thereby supporting their future use under ocean acidification conditions. Other responsive pathways harboring genes with high PC values suggest that they could be good indicators of cellular physiology although certain major pathways (e.g., photosynthesis) contain both environmentally responsive (e.g., light reactions) and unresponsive (e.g., photosystem I) genes. Hence, it is important to consider the functions of different components that make up larger biochemical pathways when investigating energy and/or nutrient flux through the cell. It is as equally important to consider the nature of the environmental conditions being tested. As discussed above, environmentally responsive iron stress genes that form complexes with high-inventory photosystems (e.g. IsiA) in iron limitation can yield high RNA-protein correlations across treatments, which may not be the case if another set of conditions were tested that lacked iron stress. These data not only fill knowledge gaps relating to long-term, nutrient-limited mRNA-protein dynamics, but also to differences and commonalities among molecular metabolisms of interrelated single- and co-limited physiologies under increasing CO_2_. A model of transcript and protein production and decay was able to reproduce correlations (in fold change units) observed for environmentally responsive genes with small initial inventories, consistent with the observations, and demonstrating the mechanics of how both transcripts and proteins can be useful in assessing ecosystem and biogeochemical function. Although these data highlight molecular dynamics fueling important biogeochemical processes, future studies examining mRNA-protein correlations over longer temporal periods (e.g., diel to several cell cycles) will help to elucidate transcriptional and corresponding protein synthesis times in the face of environmental change.

## Acknowledgements

This work was supported by US National Science Foundation Grants OCE 1851222 and OCE 1657757 and (to D.A.H., E.A.W., and F.-X.F.), OCE1924554 and OCE1850719, G.B Moore Foundation, and NIH R01 GM135709 grants (to M.A.S.).

## Supplementary Information

Supplementary Figures S1 and S2 show the experimental design and Pearson correlation coefficients, respectively. Supplemental Movie M1 provides a time course of model results transcript and protein inventories and the influence on log2 fold change results. Supplemental Dataset 1 provides the combined transcriptome-proteome dataset as an Excel spreadsheet. Tabs within the sheet include normalized biological duplicate transcript counts, biological triplicate protein spectral counts, the resulting log2 fold change dataset for transcripts and proteins, and a key for gene annotations.

## Availability of Data and Materials

All RNA-Seq data used in this study have been deposited as raw fastq files in NCBI’s Gene Expression Omnibus ^47^ and are accessible through GEO Series accession number GSE94951. All protein spectral count data used in the above analyses can be found in Walworth et al. 2016 ^23^ in Supplementary Data 4 (DOI: 10.1038/ncomms12081). Combined transcriptome-proteome dataset and gene annotations are provided as a Supplemental Dataset 1. Raw mass spectra datasets used in the above analyses are available at ProteomeXchange under the identifier PXD010515 (<u>DOI: 10.6019/PXD010515</u>). Code for the RNA-Protein model is available on GitHub under M.A.S’s repository: https://github.com/maksaito/RNA-Protein_MATLAB_model.

### Author contributions

F.-X.F., D.A.H., M.A.S. and E.A.W. designed research; N.G.W., M.D.L., M.A.S., D.M.M., M.R.M. and F.-X.F. performed research; N.G.W., M.D.L., F.-X.F., D.A.H., and E.A.W. analyzed data; M.A.S. developed the RNA-Protein model, and N.G.W., M.A.S., M.D.L., R.M. K., F.-X.F., D.A.H., and E.A.W. wrote the paper.

