## Supplemental Figures S1 and S2 for "Why Environmental Biomarkers Work: Transcriptome-Proteome Correlations and Modeling of Multi-Stressor Experiments in the Marine Bacterium *Trichodesmium*"

**Supplementary Information**

**Figure S1 Cell physiology and global transcriptional analysis** (a) Experimental design is displayed. (b) Nonmetric multidimensional scaling of Bray-Curtis dissimilarities calculated from global normalized transcripts among all biological replicates of all treatments. (c) Hierarchical clustering of Bray-Curtis dissimilarities with multiscale bootstrap resampling calculated from global normalized transcripts. Numbers at dendrogram nodes are approximately unbiased p-values calculated from multiscale bootstrap resampling.

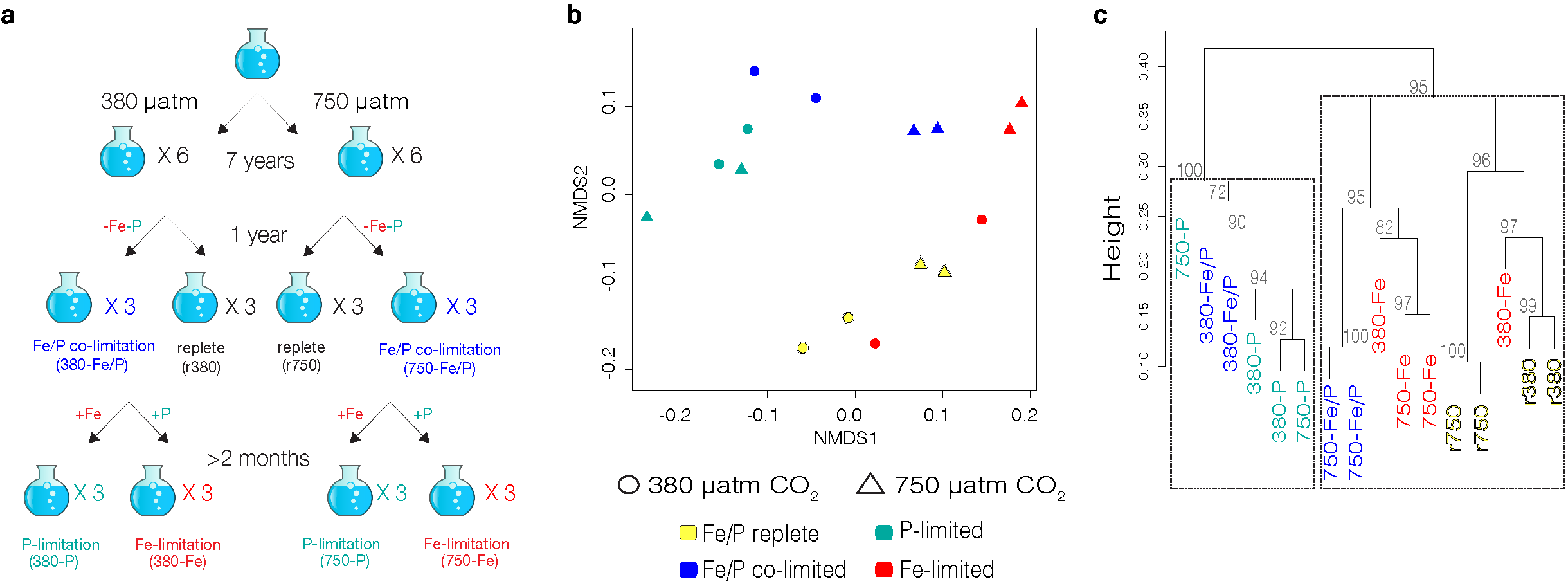

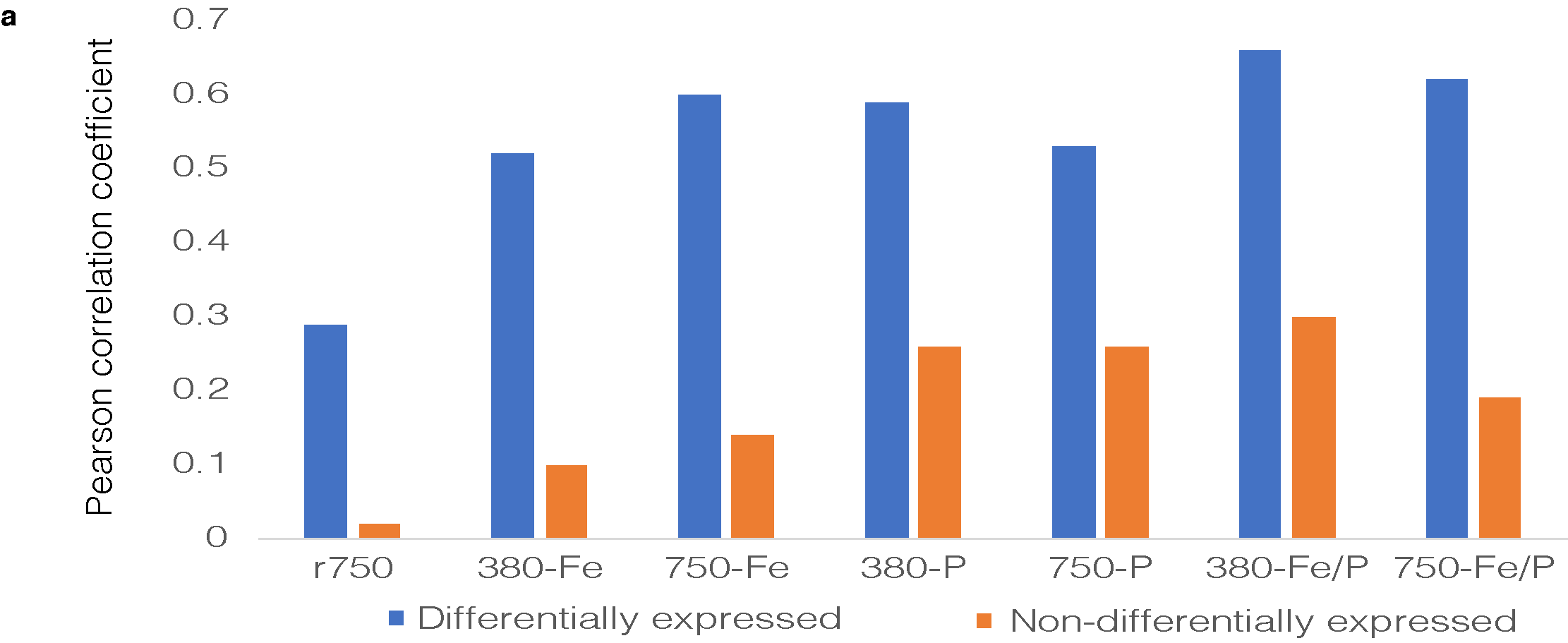

Fig. S2 **Pearson correlation coefficients of non-differentially expressed (NDE) and differentially expressed (DE) genes relative to nutrient replete 380 µatm CO_2_** Shown are average Pearson correlation coefficients per treatment of NDE (orange) and DE genes relative to r380 (blue), respectively.
